# A bioinformatic approach for the prediction and functional classification of *Toxoplasma gondii* long non-coding RNAs

**DOI:** 10.1101/2024.05.30.596643

**Authors:** Laura Vanagas, Constanza Cristaldi, Gino La Bella, Agustina Ganuza, Sergio O. Angel, Andrés M. Alonso

## Abstract

Long non-coding RNAs (lncRNAs) have emerged as significant players in diverse cellular processes, including cell differentiation. Advancements in computational methodologies have facilitated the prediction of lncRNA functions, enabling insights even in non-model organisms like pathogenic parasites, in roles such as parasite development, antigenic variation, and epigenetics. In this work, we focus on the apicomplexan *Toxoplasma gondii* differentiation process, where the infective stage, tachyzoite, can develop into the cysted stage, bradyzoite, under stress conditions. Using a publicly available transcriptome dataset, we predicted lncRNA sequences associated with this differentiation process. Notably, a substantial proportion of these predicted lncRNAs exhibited stage-specific expression, particularly at the bradyzoite stage. Furthermore, co-expression patterns between coding transcripts and lncRNAs suggest their involvement in shared processes, such as bradyzoite development. TglncRNA loci analysis revealed their potential influence on the expression of nearby coding genes, including subtelomeric genes unique to the *T. gondii* genome. Finally, with a k-mer analysis approach, we identified functional relationships between characterized lncRNAs from model organisms like *Homo sapiens* and predicted *T. gondii* lncRNAs. Our perspective led to the identification of a *T. gondii* lncRNA potentially mediating DNA damage repair pathways, shedding light on the adaptive mechanisms of *T. gondii* in response to stress conditions.

## Introduction

Transcriptomic studies have revealed that transcriptional machinery in eukaryotic organisms is capable of producing a repertoire of RNAs with little or non coding potential initially considered as transcriptional noise^1^. Over time, those non-coding RNAs exceeding 200 nucleotides were categorized as long non-coding RNAs (lncRNA) and have been proposed to carry out diverse functions in complex organisms that include regulation of cell differentiation and development, stress responses, diseases conditions and resistance to drugs^2^. At the molecular level, lncRNA’s regulatory mechanisms have been classified as *cis* acting when regulating local transcription of those genes close to a lncRNA locus. The best example of a *cis*-acting lncRNA is the X-inactive specific transcript XIST^3^. On the other hand, lncRNAs may also display its regulatory mechanism in *trans*. This process depends on the production of transcripts that exert their function by interacting with proteins, DNA or RNA molecules^4^. Comparative analysis has revealed that lncRNA sequences are not well conserved across species, making lncRNAs functional prediction a difficult task^5^. Nevertheless, new computational approaches have been applied and functional related lncRNAs can now be predicted by *k*-mer content, co-expression networks and secondary structure homology prediction^6–8^. Additionally, the prediction of novel lncRNAs in non-model organisms, such as pathogenic parasites, could be now feasible through computational analysis like those mentioned above, making a promising starting point in the field. In the context of parasitology, lncRNA studies have been principally focused on the host-pathogen interaction process. However, there are a few studies documenting the lncRNA role at parasite development, antigenic variation and epigenetics^9,10^. Parasite development is a complex process, crucial for organism adaptation to its environment, and governed by gene regulation. This has underscored the relevance of studying lncRNAs in this context. In fact, a documented lncRNA has been identified as a regulator of the differentiation process in both *P. falciparum* and *T. brucei*^11,12^, and it has also been implicated in the sexual development of helminths^13^. Additionally, a lncRNA named Tg-ncRNA-1 was proposed as regulator of Toxoplasma gondii development^14^. The apicomplexan *T. gondii* is an obligate intracellular parasite with a complex life cycle that compresses definitive hosts (felines) where parasites go through a sexual development and intermediate hosts (not only mammals and birds) where the asexual phase of the parasite takes place. Within the intermediate hosts, the parasite adopts an actively infective form that colonizes cells, replicating at high rates, the tachyzoite, until differentiation to a latent bradyzoite which encysts in different tissues^15^. Transition of tachyzoite to bradyzoite stage is a cell differentiation process coordinated and regulated by differential expression of key regulators. *In vivo*, this is induced by a stress context provoked by the immune response of the host, but this process can also be stimulated by physical and chemical agents *in vitro*^16,17^. It has been established by a great variety of studies that model organisms express lncRNAs as a response to stress conditions like DNA damage, hypoxia and heat-shock^18–20^. Considering the differentiation process in *T. gondii* as a stress response which until now has been actively faced from the genome coding point of view, analyzing the lncRNAs role can contribute to understanding the plasticity of non-coding genome of *T. gondii* on adaptation and response to stress conditions as well as other biological situations. In this work we explored a publicly available transcriptome dataset focused on the *T. gondii* tachyzoite to bradyzoite differentiation process at the TgME49 strain with the aim of predicting lncRNAs sequences and their functional association. We could observe that a great proportion of the predicted lncRNAs for the ME49 strain were specifically expressed at the bradyzoite stage. Additionally, clustering methodologies helped us to infer that co-expressed coding transcripts and lncRNAs could share participation on the same processes, like those necessary for bradyzoite development. Besides, our study of the TglncRNA loci revealed their possible influence on the expression of nearby coding genes. Specifically, we could observe by this analysis the potential influence on subtelomeric genes like TgFAM, a family of genes uniquely enriched at *T. gondii* genome^21^. Finally, we implemented a *k*-mer analysis perspective^6^ for determining functional relationships between characterized lncRNAs from a complex organism like *Homo sapiens* and the predicted *T. gondii* lncRNAs (TglncRNAs). This approach led us to identify one TglncRNA potentially mediating the DNA damage repair pathways in *T. gondii*. In summary, our study underscores the significance of TglncRNAs in the regulatory landscape of *T. gondii*, offering novel insights into its adaptive strategies and potential therapeutic targets.

## Results

### Prediction of *Toxoplasma gondii* long non-coding RNAs

Since there is limited research on *T. gondii* lncRNAs, we designed a bioinformatics pipeline with the aim of predicting novel TglncRNAs from available RNAseq data (Fig. 1a). We focused our analysis at the stage transition of tachyzoite to bradyzoite since this cell differentiation was previously documented as probably regulated by a TglncRNA^14^. In this sense we intended to expand the repertoire of TglncRNAs potentially expressed along the stage transition. To accomplish our aim, we employed a validated transcriptomics data set for the tachyzoite to bradyzoite stage conversion in the ME49 strain^22^. Next we assembled and quantified a merged transcriptome for the two stages, with focus on new and unannotated sequences to finally classify these as coding or non-coding as was described in the material and methods section. Those sequences predicted as non-coding were filtered by sequence length > 200 nt as was documented in previous studies^10^ and accepted those transcripts as predicted TglncRNA for *T. gondii*; as result a total of 656 sequences were predicted as TglncRNAs. Additionally 320 of those transcripts (∼ 49%) were validated by a blastn analysis against expressed sequences tag evidence (ESTs) for *T. gondii*. Genome localization and relevant data for TglncRNA sequences are available at Supplementary Table S1. Then we explored the characteristics associated with the predicted TglncRNAs. Initially, we noted that TglncRNAs exhibited lower expression levels compared to coding transcripts (Fig. 1b). In addition, TglncRNAs were relatively shorter and had a lower percentage of GC in comparison with coding transcripts (Fig. 1c). Next, we wondered if the predicted TglncRNAs codified for a protein product. In this sense, we searched for homologs at the uniprot database^23^ for the predicted sequences, finding that only 21 of them are related with at least one uniprot reference sequence (Supplementary Table S1). Among these, 12 were related with 2 *T. gondii* reference sequences: a Citochrome b protein (O20672) and a SAG-related protein (P13664; *E* −*value <* 0.001). We have not excluded these sequences for further analysis since it is known that non-coding RNAs may act as bifunctional RNAs^24^.

**Figure 1.**
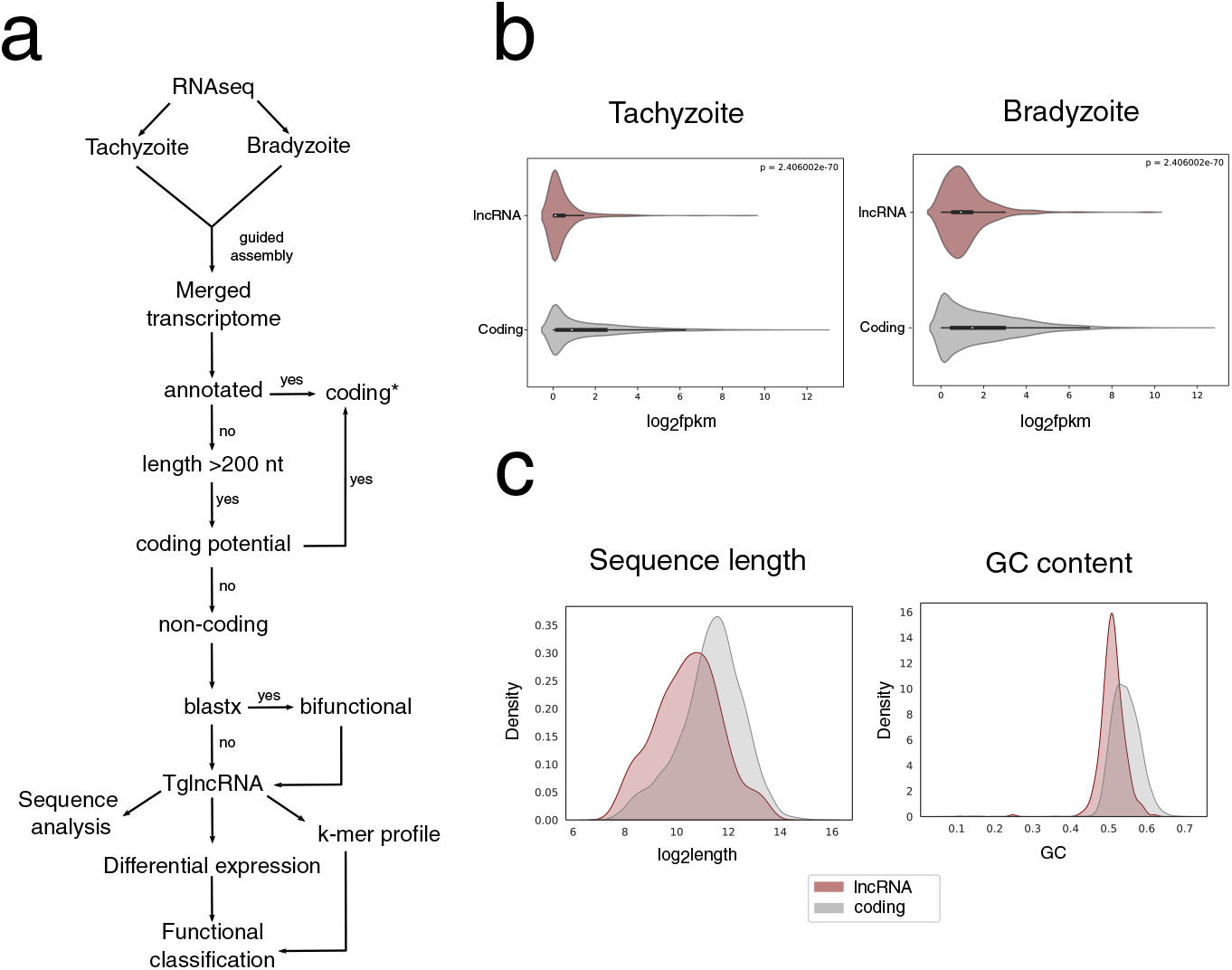
Bioinformatic pipeline and sequence properties of the predicted TglncRNAs. (a) Schematic representation of the bioinformatic approach for TglncRNAs prediction and analysis implemented in this work. *= Noncoding annotated transcripts, like rRNA and tRNA, were also filtered from transcriptome analysis. (b) Relative mean expression comparison for TglncRNAs and coding transcripts at tachyzoite and bradyzoite (*p*-value<0.001); (c) Density histogram. Upper panel: sequence length distribution for predicted TglncRNAs and coding transcripts; sequence lengths were *log*_2_ transformed. Bottom panel: Total GC content distribution for TglncRNAs and coding transcripts.

We also studied the TglncRNAs sequence distribution. As shown in Figure 2a, predicted sequences are distributed along the reference chromosomes of the ME49 strain. Additionally we mapped the relative position of the predicted TglncRNA loci to protein-coding genes. As results we observed that a great proportion of the loci are close to promoter regions (74%) while the rest of the loci were classified as intergenic (Fig. 2b).

**Figure 2.**
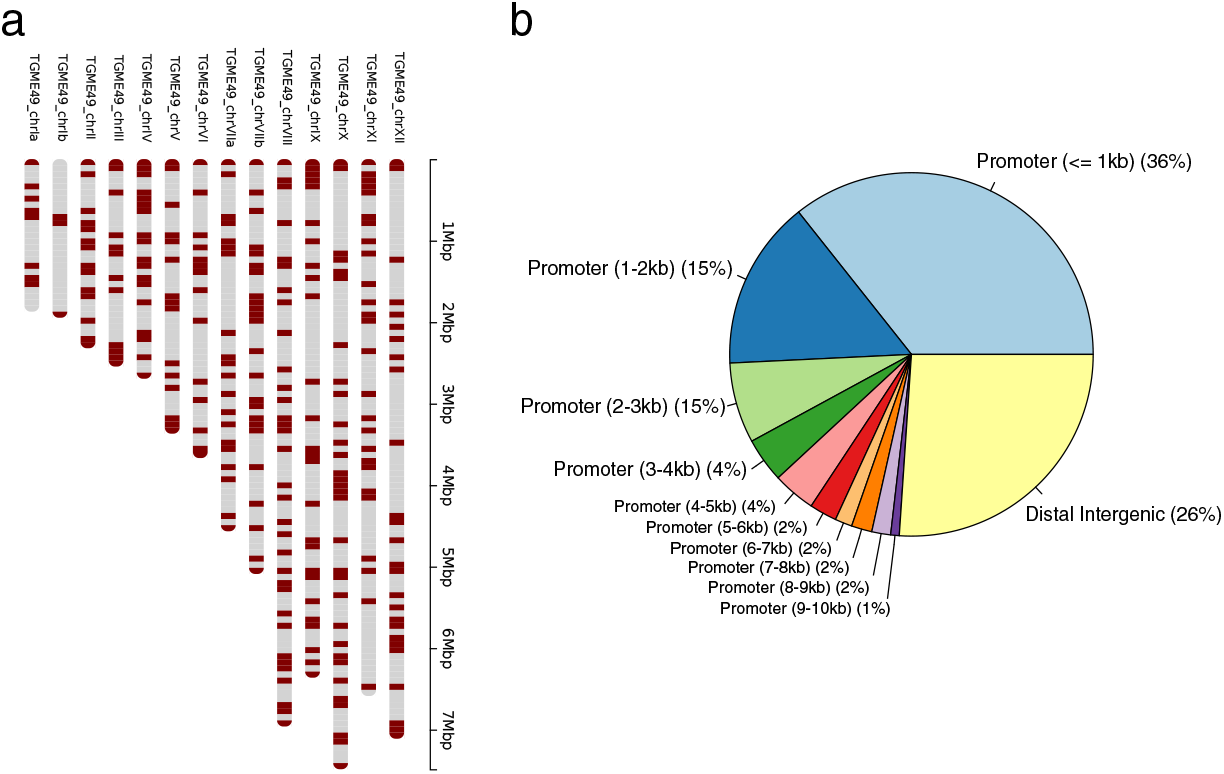
Genomic location of predicted TglncRNAs. (a) Distribution of predicted TglncRNAs along *T. gondii* chromosomes. Each colored band represents a TglncRNA locus and chromosome length scale is expressed in mega bases pairs (Mbp); (b) Pie plot that summarizes the percent of TglncRNA loci relative to coding gene positions. Distance to coding gene promoters is expressed in kilobases (kb). Distal intergenic are those more than 10kb away from the gene promoter.

### Comparative analysis between TglncRNAs from VEG and ME49 strains

Since previous studies over *T. gondii* had predicted a repertoire of TglncRNA for the VEG strain, we conducted a comparative analysis using the results from the ME49 strain. Summary of our observations, shows that the explored characteristics of the TglncRNA sequences exhibit similarities across studies (Table 1). From the transcriptomics point of view, 206 sequences predicted by our analysis have homology with the previously documented VEG TglncRNAs^25^ (Fig. 3A). This observation is interesting, since a higher similarity could be expected between strains sequences. However, it has been documented that lncRNAs expression is strain specific in other model organisms like *Drosophila*^26^. Although a great proportion of the predicted ME49 TglncRNAs were not detected over the transcriptomics of VEG strain, genomic comparative analysis revealed that 645 from 656 sequences are present at VEG strain, mostly with high synteny, revealing a high level of conservation at least from genomics point of view (Fig. 3B).

**Table 1.** Summary table for TglncRNAs principal characteristics at ME49 and VEG strains. Total: total number of analyzed sequences; Max length: maximum sequence length; Mean length: mean length of analyzed sequences; GC%: GC percent of sequences; ETSs: number of sequences with at least one match for expressed sequences tags (ESTs).

|  | ME49 | VEG | Coding(ME49) |
| --- | --- | --- | --- |
| **Total** | 656 | 889 | 8322 |
| **Max length** | 12141 | 15541 | 52319 |
| **Mean length** | 2042 | 1640 | 3530 |
| **Min length** | 201 | 211 | 114 |
| **GC%** | 51,00 % | 51,00 % | 54,00 % |
| **ESTs** | 320 | 531 | – |

**Figure 3.**
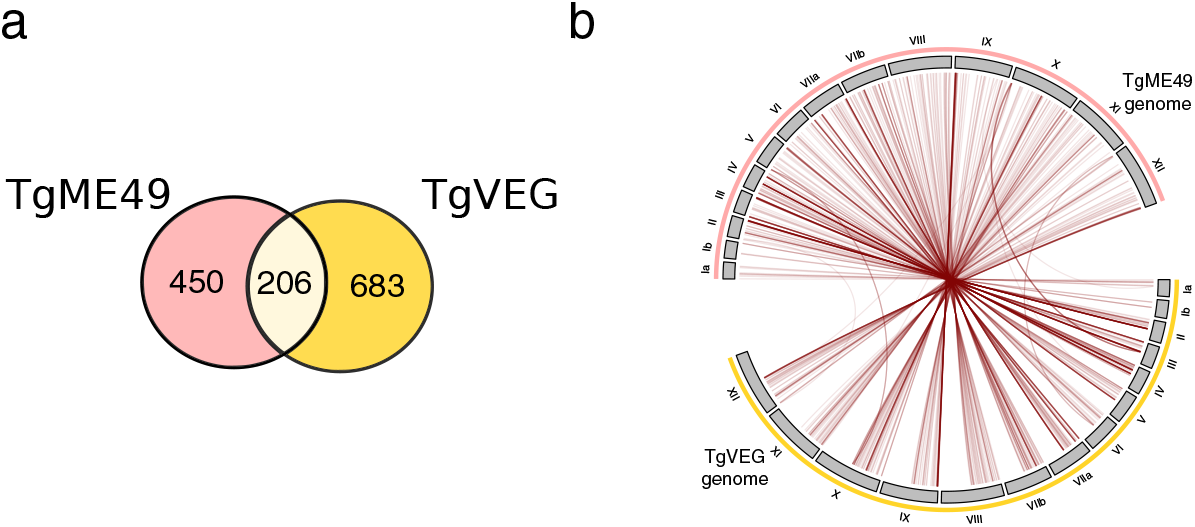
Comparative analysis of *T. gondii* lncRNAs between ME49 and VEG strains. (a) Venn diagram representing shared sequences of TglncRNAs between ME49 and VEG transcriptomes. (b) Circos plot that shows syntenic positions for 645 TglncRNAs from ME49 strain at VEG strains.

### Differential Expression of predicted lncRNAs during stage conversion

Since early studies proposed a role for lncRNAs in cell differentiation^27^ we wondered if a differential expression of our predicted TglncRNAs would be observed during tachyzoite to bradyzoite transition. Based on that, we performed a differential expression analysis over the assembled transcriptome, with focus on non-coding transcripts. As result, we determined 217 differentially expressed lncRNAs. Three of these were significantly upregulated in tachyzoite state while the rest, 214 were significantly upregulated in bradyzoite state (*pad j <* 0.05 and *log*_2_ *f c >* 2, Fig 4A and Supplementary Table S2). It is worth mentioning that a previous work in *T. gondii* argues that a TglncRNA, Tg-ncRNA-1 regulates bradyzoite formation^14^. The locus from that transcript is located at *TGME*49_*chrVI* : 66, 556..69, 156 where we were able to determine one sequence poorly expressed at tachyzoite and bradyzoite stages. In our work, that sequence was not differentially detected at the bradyzoite stage (*MSTRG*.3577.1, Supplementary Table S1). We highlight one TglncRNA, *MSTRG*.6814.2, since it was detected as differentially expressed at the bradyzoite stage with high confidence (*pad j* = 1*e* − 89). Additionally, our analysis detected *MSTRG*.6814.2 (from now TglncRNA-5J) as the most expressed TglncRNA at the bradyzoite stage. Under this context, we proposed to validate TglncRNA-5J expression in addition with other highly expressed TglncRNAs at the bradyzoite stage.

**Figure 4.**
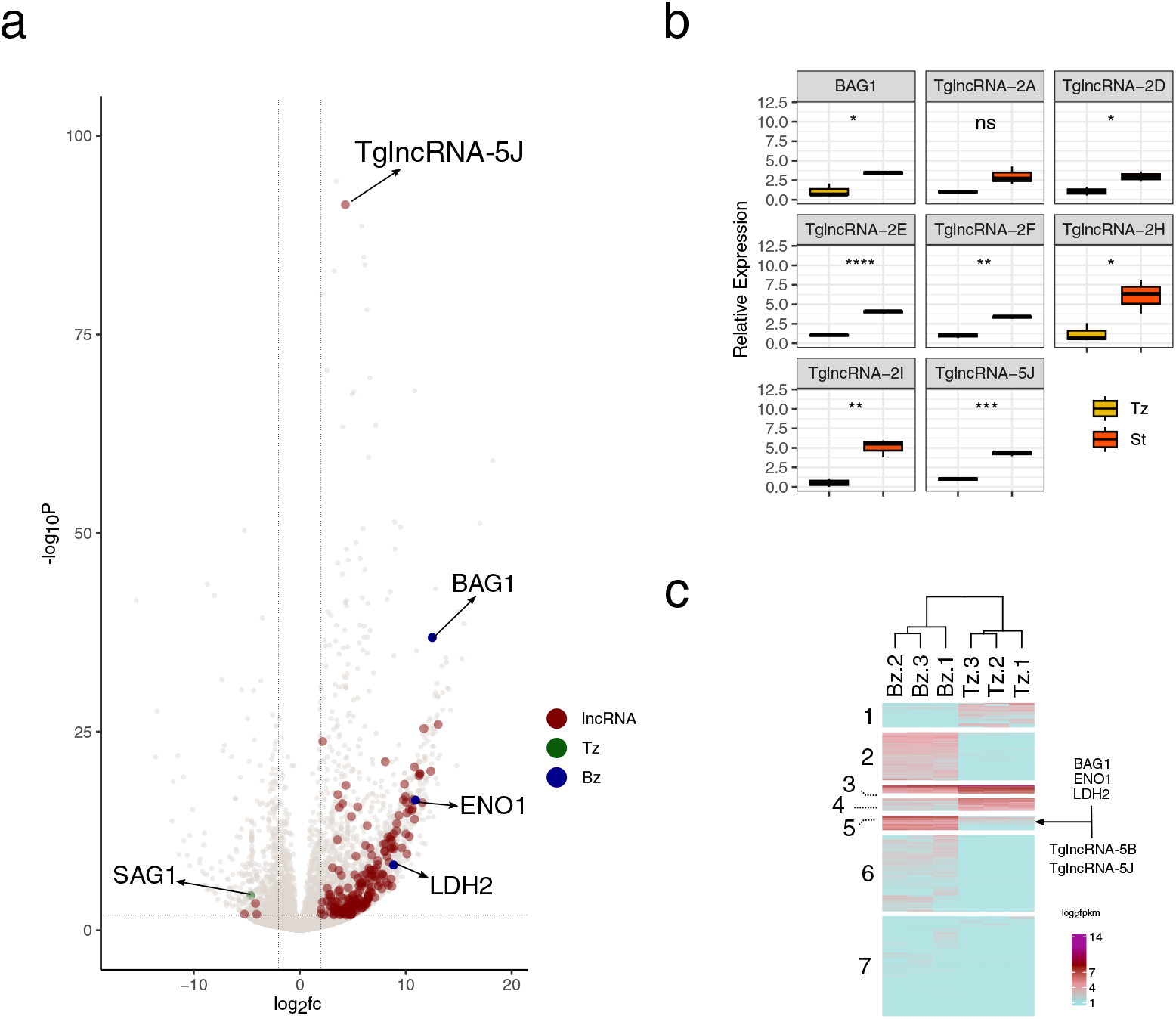
TglncRNAs differential expression between tachyzoite and bradyzoite stages. (a) Volcano plot for differentially expressed transcripts between tachyzoite and bradyzoite stages. Differentially expressed lncRNAs (*pad j <* 0.05 & *log*_2_ *f c >* 2) are highlighted (red). The most relatively expressed TglncRNA at the bradyzoite stage is highlighted (TglncRNA-5J = *MSTRG*.6814.2). Standard transcripts normally expressed for each stage are also highlighted. Tachyzoite (green): surface antigen-1 (SAG1). Bradyzoite (blue): bradyzoite antigen (BAG1), enolase-1 (ENO1), lactate dehydrogenase-1 (LDH1). Axises represent the logarithmic transformation of the fold change (*log*_2_ *f c*) and the minus logarithmic transformation of the adjusted p-values (−*log*_10_*P*; Benjamini and Hochberg). (b) qPCR analysis for 8 selected TglncRNA transcripts. Tz: tachyzoite control; St: heat-shock stressed extracellular tachyzoites. ns: *p >* 0.05, ∗ : *p <*= 0.05; ∗∗ : *p <*= 0.01, ∗∗∗ : *p <*= 0.001 and ****: *p <*= 0.0001. (c) Heatmap plot for differentially expressed transcripts, coding and lncRNA transcripts. The heatmap was sectioned in 7 transcript clusters. Standard transcripts normally expressed for the bradyzoite stage are highlighted at cluster 5 grouped with the two most highly expressed lncRNA transcripts (TglncRNA-5B and TglncRNA-5J). Bar color scales represent the logarithmic transformation of the relative expression (*log*_2_ *f pkm*; fragment per kilobase millon).

### Stress induced TglncRNA expression in extracellular tachyzoites

During the lytic cycle the *T. gondii* tachyzoite exits the cell to the extracellular space where it must quickly find another nucleated cell to invade and continue with the replicative process. The passage to the extracellular environment represents a condition of environmental stress in itself that accompanies changes in gene expression. Interestingly, the application of some stress situations in the extracellular tachyzoite affects the expression of many genes, including some that are specific to the bradyzoite stage^28^. In this context we reasoned that the differential expression of *T. gondii* lncRNA transcripts at stress contexts like those that trigger tachyzoite to bradyzoite conversion, could be mimicked by stressing extracellular tachyzoites. With such a simple experiment we aimed to validate the expression of lncRNAs predicted by our analysis. Taking into account relative expression values from our transcriptomics analysis (Fragments Per Kilobase Million; fpkm) we selected the 10 most abundant TglncRNAs from our differential analysis at bradyzoite stage and designed qPCR primers to quantify TglncRNA transcript expression with aim of validate our predictions in a stress induced experiment. Selected TglncRNA for our analysis are listed at Table 2. Extracellular tachyzoites were incubated under heat-shock conditions for 2 hours with the aim of inducing TglncRNA transcript expression to be quantified by qPCR. As can be seen at Figure 4b we were able to detect amplification of 7 of the 10 selected TglncRNAs. From the 7 amplificated TglncRNA we confirmed that heat-shock stress induced differential expression on 6 TglncRNAS (*p*−*value <* 0.05). Additionally, we highlight that the most upregulated TglncRNA (TglncRNA-5J) proposed by our predictions was the most relatively expressed transcript in our experimental validation, an observation that prompts us to hypothesize that TglncRNA-5J could have a role under stress conditions. In conclusion, we confirmed that putative TglncRNAs of *T. gondii* could be expressed under stress conditions in extracellular tachyzoites. To note, the bradyzoite marker gene BAG1 also accompanies the upregulation of the TglncRNAs, opening the option that these could also be upregulated during the conversion to bradyzoite.

**Table 2.** Selected TglncRNA transcripts for qPCR experiment. Values indicates bradyzoite (Bz) and tachyzoite (Tz) relative abundance (*log*_2_ *f pkm*) detected from transcriptomics. Cluster: the cluster number where the TglncRNA is founded in the heatmap representation (Fig. 4b).

| Name | lncRNA ID | Fold change<br>(Bz/Tz) | Relative abundance (log2fpkm) |  | Cluster |
| --- | --- | --- | --- | --- | --- |
|  |  |  | Stress | Control |  |
| TglnRNA-2A | MSTRG.1940.1 | 6.43 | 3.12 | 0.05 | 2 |
| TglnRNA-5B | MSTRG.4604.1 | 3.08 | 4.46 | 1.56 | 5 |
| TglnRNA-2C | MSTRG.3554.1 | 7.09 | 4.31 | 0.04 | 2 |
| TglnRNA-2D | MSTRG.4702.1 | 2.16 | 3.95 | 1.79 | 2 |
| TglnRNA-2E | MSTRG.1204.1 | 2.59 | 3.83 | 1.38 | 2 |
| TglnRNA-2F | MSTRG.6333.1 | 3.59 | 3.81 | 0.86 | 2 |
| TglnRNA-2G | MSTRG.1941.1 | 4.36 | 3.52 | 0.47 | 2 |
| TglnRNA-2H | MSTRG.6839.1 | 12.36 | 3.25 | 0 | 2 |

### Functional analysis of differentially expressed Tg-lncRNAs

Taking in consideration that the lncRNAs function prediction by sequence homology is difficult, we reasoned that co-expressed genes with known function would be useful to associate the predicted TglncRNAs to a common enriched process or function. In accordance, we focused on the annotated gene transcripts that were differentially expressed at tachyzoite to bradyzoite transition and clustered them using their relative expression (*log*_2_ *f pkm*) between parasite stages. Following this analysis, seven clusters emerged (Fig 4C), with the predicted TglncRNAs distributed among five of them (Supplementary Table S3). Clusters 7 and 6 contain a great proportion of lncRNAs,110 and 81 respectively, while cluster 1, 2 and 5 contain the smallest number of TglncRNAs, and clusters 3 and 4 none. By this method the different TglncRNAs can be grouped on the basis of its co-expression with genes of different biological processes (BP) and molecular function (MF). In particular, TglncRNAs in cluster 7 could be related with the cytoskeletal and cell motility (BP) and lyase activity (MF). TglncRNAs of cluster 6 could be related to microtubule-based movement (BP) and peptidase activity (MF). In Cluster 2, 22 lncRNAs were coexpressed with transcripts related with peroxisome organization (BP) and acyltransferase activity (MF). Cluster 1 was poorly integrated by TglncRNA transcripts, finding only 2, and was related to intracellular protein transport (BP) and acyltransferase activity (MF). A separate item is cluster 5, which was integrated only by 2 TglncRNAs (named TglncRNA-5J and TglncRNA-5B). Carbohydrate metabolic processes are the most enriched BP and oxidoreductase activity MF. A close inspection over this cluster led us to find that it is composed of transcripts that codify for relevant proteins in the bradyzoite stage like BAG1, ENO1, LDH2, CST7, MAG2 and PMA1^29–32^. In fact, we observed the presence of BFD2, recently documented as a RNA binding protein that regulates BFD1, a key regulator for bradyzoite development^33^. This last observation allows us to hypothesize, once again, that these TglncRNA transcripts may have a relevant role in bradyzoite development in conjunction with the other members of cluster 5. In conclusion, the co-expression of certain TglncRNAs with a cluster of genes could be indicating some participation of some of those TglncRNAs in such molecular and biological processes. Of course, a great proportion of coding genes with specific function or unknown function at *T. gondii* (hypothetical, subtelomeric or pathogenic factors) have no mapped GO ontologies which difficult the association of specific ontologies to TglncRNAs. In this sense we explored functional analysis by loci genomic distance of TglncRNA to coding genes.

### *cis*- and *trans*-acting putative associated mechanism of differentially expressed Tg-lncRNAs

The functions of TglncRNAs can be grouped based on whether their target of action is nearby (*cis*-acting), or distant (*trans*-acting) from the lncRNA loci^34^. We hypothesized that a predicted TglncRNA locus close to a protein-coding gene would affect gene expression in cis. Our observations revealed that 71% of the differentially expressed lncRNA transcripts (153) at the bradyzoite stage were classified as closer to promoter regions of T. gondii genes (Fig. 2b). A total of 222 genes could be influenced by these close TglncRNA loci and 46 of these 222 genes were differentially expressed between *T. gondii* stages (Supplementary Table S4; predicted *cis*-TglncRNAs). Five genes of the 46 above mentioned, were downregulated at the bradyzoite stage: three are annotated as hypothetical proteins (*TGME*49_289910, *TGME*49_249300 and *TGME*49_278780), one as tachyzoite ROP2A (also known as ROP24,*TGME*49_215785)^35^ and one as a *Toxoplasma gondii* family C protein (FamC, *TGME*49_200130), a putative subtelomeric gene. In fact The locus for the TglncRNA-5J (TglncRNA up-regulated at bradyzoite stage) is located upstream (*∼* 2.5*kb*) of the ROP2A locus and it is a good candidate as *cis*-acting regulator of ROP2A gene expression (Fig. 5a). We noted that a hypothetical protein (*TGME*49_275310) is close to TglncRNA-5J locus, but it was not considered since it was not detected by our differential expression analysis at the tachyzoite or bradyzoite stages. Forty-one protein coding transcripts were found upregulated at the bradyzoite stage highlighting the presence of the BAG1 gene (*pad j <* 0.05 & *log*_2_ *f c >* 2). Although a significant proportion of these genes encodes for uncharacterized proteins (24) it is important to highlight that we observed 5 putative subtelomeric genes as differentially expressed (Fig 5B and Supplementary Table S4). In this context, cis-acting TglncRNAs in close proximity to subtelomeric genes implies their potential role in regulating the expression of a diverse array of subtelomeric genes. In fact this role of lncRNAs at subtelomeric gene regulation was previously observed in *Plasmodium* at var genes regulation^36^. We next focused on those TglncRNAs loci classified as distal intergenic in this work and differentially expressed at bradyzoite stage. We reasoned that these TglncRNAs would act in trans^4^. *trans*-acting mechanism of lncRNAs involves their interaction with other molecular entities, such as coding transcripts. This interaction can regulate the translation, mRNA stability or splicing^37–40^. In accordance with this, we performed a computational analysis with the aim of predicting RNA-RNA interactions between up-regulated TglncRNA sequences and differentially expressed coding transcripts by taking into account RNA secondary structure (RIBlast^41^). Our analysis revealed that 43 TglncRNAs have a total of 117 coding transcript targets at the bradyzoite stage (Supplementary Table S4; ; predicted *trans*-TglncRNAs). Among these transcripts, a majority of hypothetical genes are identified but also several transcription factors of the TgAp2 family. Our hypothesis is that co-expression of the TglncRNAs and their targets is needed for RNA-RNA interaction, which gives us additional evidence for the predicted interaction by RIBlast. In this sense we calculated correlations between the 43 TglncRNA and their predicted targets based on relative expression to construct a correlation network. In this way, any strong relationship between the TglncRNAs and their targets could be revealed. Under these context links of the network were represented by the calculated correlations using relative expression of both coding and non-conding transcripts while all the transcripts were considered the nodes of the network. Additionally we performed a community analysis over the correlation network with the aim of detecting those highly interconnected nodes. In the correlation network (Fig. 5c) the high level of relationship between the TglncRNAs and their predicted RIBlast targets may be observed (1347 links between transcripts; *r* −*value >* 0.6 & *p*−*value <* 0.01). Additionally 4 communities of highly interconnected nodes were observed; community composition is listed in Supplementary Table S4 (Predicted *trans*-TglncRNAs). Since a network community is a subset of nodes that are highly interconnected with each other in comparison with the rest of the nodes in the network^42^, we reasoned that communities should be composed by TglnRNAs and coding transcripts that show high correlation and a predicted RIBlast interaction with each. In fact when we explored the communities we realized that the above hypothesis could be confirmed. As an example we observed that coding transcripts for a SAG-related protein (SRS17A), argonaute (AGO) and DICER integrate the same community with at least one of the predicted interacting TglncRNAs (i.e: *MSTRG*.5823.7 and *MSTRG*.5823.10 integrate the same community as SRS17A. Supplementary Table S4; predicted *trans*-TglncRNAs). In community 2 the presence of an apiAP2 transcription factor and a DNA methyltransferase (DNMTb) is highlighted; both associated to gene and chromatin regulation. It is important to mention that communities 3 and 4 are integrated with a great proportion of transcripts that encode for hypothetical proteins, and at least one of the TglncRNAs that was predicted to interact with those coding transcripts share the same community. With relation to the Focusing in analyzed TglncRNAs by qPCR in this work, only one interaction was predicted by RIBlast and was for *MSTRG*.1940.1 (Tg-lncRNA-2A,Fig. 5c) with a Kazal serine protease inhibitor. We highlight that both share community 1 (Supplementary Table S4; predicted *trans*-TglncRNAs). While these last findings do not conclusively confirm a physical interaction between these RNA molecules, they underscore the necessity for further experimental studies in this area. These observations warrant exploration to validate their significance.

**Figure 5.**
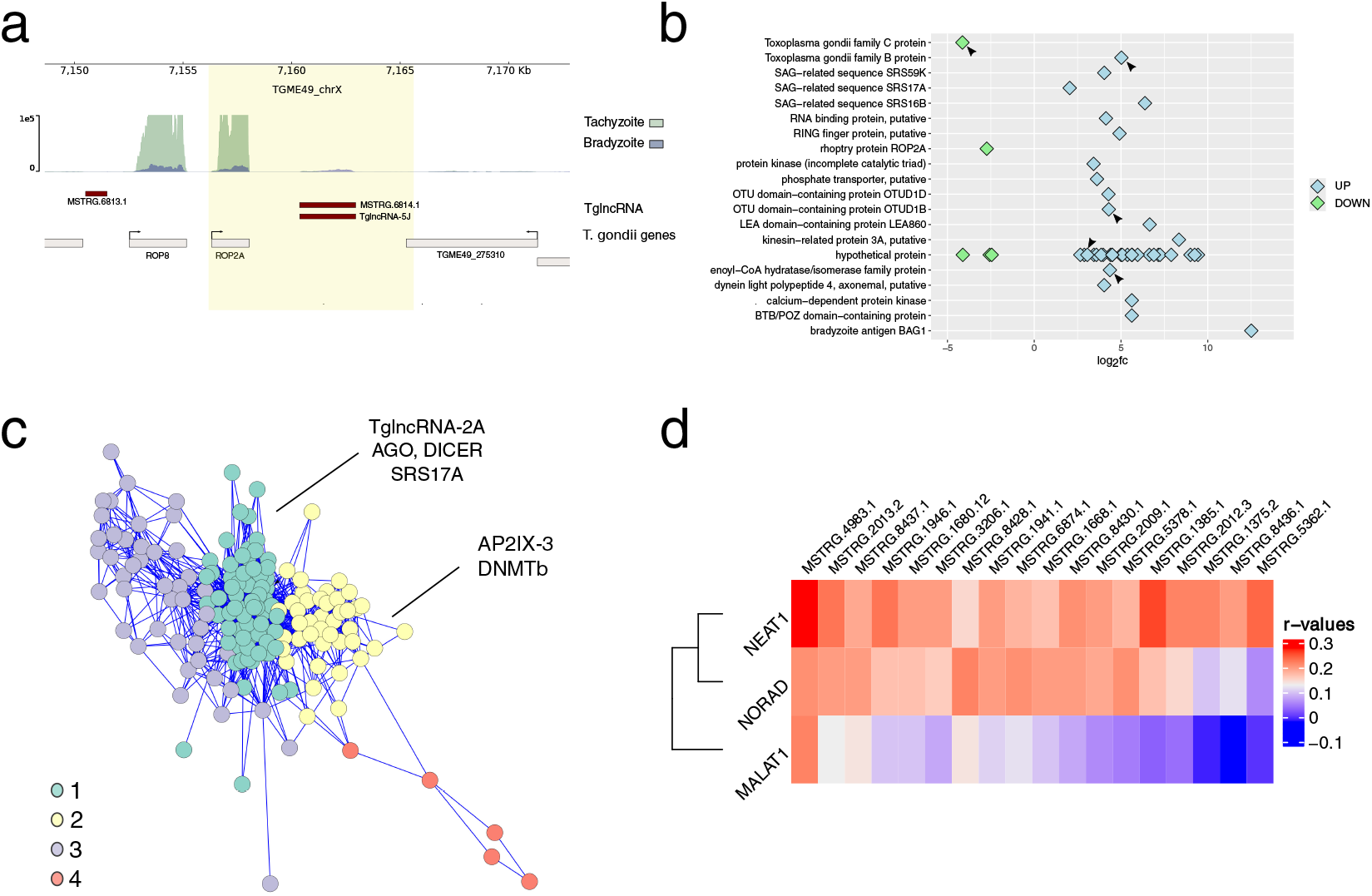
Functional annotation of predicted TglncRNAs. (a) Genomic localization of TglncRNA-5J at chromosome *TGME*49_*chrX* : 7, 148, 623 − 7, 172, 814. Tachyzoite and Bradyzoite RNAseq coverage. TglncRNA: genomic position of lncRNA loci. *T. gondii* genes: genomic position of coding genes loci. (b) A Point plot for *log*_2_ *f c* of differentially expressed genes at the bradyzoite stage. The plotted *f c* corresponds to genes near to TglncRNAs loci. Points are highlighted as follows: light blue upregulated (UP) and light green downregulated (DOWN). Arrow heads highlight subtelomeric genes: OTUD1B (*TGME*49_237894), FamB (*TGME*49_279460), FamC (*TGME*49_200130), hypothetical protein (*TGME*49_274280), enoyl-CoA (*TGME*49_317705). (c) Correlation network composed by TglncRNAs and mRNA differentially expressed at bradyzoite stage and predicted by RIBlast as TglncRNAs interactors. More interconnected nodes are grouped in communities (1,2,3,4) and only significant links are plotted (*p*−*value <* 0.01 & *r* −*value >* 0.6). The highlighted mRNAs in communities 1 and 2 share the community with at least one of the TglncRNAs that was predicted by RIBlast as an interactor. Examples listed in the main text are highlighted. AGO: argonaute-1; DICER: ribonuclease type III Dicer; SRS17A: SAG-related sequence; AP2IX-3: AP2 domain transcription factor; DNMTb: DNA methyltransferase. (d) Heatmap plot for correlations of k-mer profiles between 18 TglncRNAs and HslncRNAs: MALAT1 (*URS*0000*A*77400_9606), NEAT1 (*URS*000075*DAEC*_9606), NORAD (*URS*000075*CEFB*_9606). Scale bar represents values for correlation pearson coefficient (*r* −*values*)

### Prediction of TglncRNA binding proteins by *k*-mer analysis

LncRNA functional annotation is still a challenge since linear sequence alignment fails to predict homology due to the lack of direct evolutionary conservation^5^. Nevertheless, studies demonstrated that lncRNAs with similar *k*-mer content have been associated with similar functions^6^. In this sense, we implemented a *k*-mer based approach that consists in determining the *k*-mer profile of each of the predicted 656 TglncRNAs. This profile was then employed to scan each position weighted matrix (PWM) from each RNA binding-protein (RBP) motif from model organisms stored at CisBP-RNA database (CisBP-RNAdb^43^). Our hypothesis was that if a PWM for a RBP motif at the CisBP-RNAdb matched over a TglncRNA sequence, then it would be possible to predict a RBP binding site for that TglncRNA and associate function to the TglncRNA based on the predicted protein interaction. We performed a *k*-mer count of *k*− 5 to *k*− 7 to then calculate scores against all the PWM in CisBP-RNAdb.

As a result we obtained a matrix of weighted scores for each PWM vs TglncRNA comparison. A distribution of weights for TglncRNA *k*-mer profiles shows that a population of high scores was observed at a *k* − 7 profile that corresponds to 18 PWMs matching along the 656 TglncRNAs (Supplementary Fig. S1). We kept the ten TglncRNAs with highest scores for the 18 PWMs.

As a result we obtained a total of 81 unique TglncRNA sequences that matched with those PWMs. These 81 TglncRNAs were employed in the subsequent analysis. In this way we were able to predict the interacting RBPs for TglncRNA sequences. It should be noted that each of the predicted RBPs has an homologous protein at the T. gondii proteome (Supplementary Table S5; predicted RBP). Next, we searched at RAIDdb (RNA association interaction database^44^) for lncRNAs that were confirmed experimentally as interactors for the RBPs predicted by our analysis. Based on that, we selected for our analysis 8 well studied lncRNAs at Homo sapiens: ANCR, MALAT1, CRNDE, BCYRN1, NEAT1, NORAD, XIST, HOTAIR. We performed a correlation analysis at the *k*-mer profile level between the selected *Homo sapiens* lncRNAs (HslncRNAs) and the 81 TglncRNAs. The obtaining correlation matrix reveals relationships between the analyzed sequences; positive correlation values (*r* −*value >*= 0.2 & *p*−*value <* 0.01) between 18 TglncRNAs and 3 HslncRNAs (NEAT1, NORAD and MALAT1) were observed (Fig 5D). Furthermore, 4 TglncRNAs (*MSTRG*.1941.1, *MSTRG*.2013.2,*MSTRG*.4983.1 and *MSTRG*.8430.1) show positive correlations with more than one of 3 HslncRNAs (Fig 5D and Supplementary Table S5; HslncRNA correlations). *Homo sapiens* NORAD (non-coding RNA activated by DNA damage) contain multiple binding sites for Pumilio proteins (PUM), a family of proteins that regulate gene expression by mRNA binding at 3’-UTR^45^. Six of the 18 TglncRNAs sequences could be considered as NORAD-like sequences since PWD motifs were detected in each sequence and *k*-mer profiles show positive correlations with *H. sapiens* NORAD (Supplementary Table S5; HslncRNA correlations).In fact, our sequence analysis reveals that repetitions in a the TgNORAD-like sequence (*MSTRG*.2013.2, renamed *TglncRNA*_*NORAD*) conserve the documented structural motifs which allows us to infer functionality for this sequence^45^ (Sumpplementary Fig. S2). When we explored the *T. gondii* annotated genome it was possible to find 2 genes that encode for homologous PUM proteins among others RBPs like NONO and FUS (Supplementary Table S5; predicted RBP). Since functional association of PUM proteins with 5’-UTR and 3’-UTR from coding transcripts was documented in *Plasmodium falciparum*^46^, we explored UTRs of coding transcripts from *T. gondii* with the aim of locating PUM binding motif. We detected 14 gene encoding transcripts with 5’-UTRs positive PUM binding motif. Furthermore, a GO enrichment analysis performed over these genes showed biological processes like DNA replication (*GO* : 0006260), DNA metabolic process (*GO* : 0006259) and DNA recombination (*GO* : 0006310; Supplementary Table S7). These GOs are related with 14 genes among which we may highlight those that participate in DNA damage repair like TgRAD51 (*TGME*49_272900) and TOPOII (*TGME*49_312230). The PUMs protein represses traduction translation of the mRNAs to which it binds. It is considered that the expression of an lncRNA with PUM motifs could be titrating the PUM protein, releasing the traductional repression of binding mRNAs^18^. In this context, we consider that the *TglncRNA*_*NORAD* could be expressed in situations of DNA damage/stress, allowing the expression of mRNAs that code for proteins of the DNA damage response. In conclusion, our data suggests that lncRNA-associated DNA damage regulation could be conserved in T. *gondii*, opening a new field of study, although further validation is needed.

## Discussion

In this work we designed a bioinformatic approach with the aim of exploring the *Toxoplasma gondii* transcriptome at tachyzoite to bradyzoite stage differentiation. We focused on those unannotated transcripts, with no or poor coding potential of length > 200 nucleotides, that are commonly known as long non-coding RNAs or lncRNAs. Our predictions reveal a total of 656 TglncRNAs detected from *T. gondii* transcriptomes. Of these, 217 were highlighted as differentially expressed between parasite stages. Interestingly, a great proportion of differentially expressed TglncRNAs were detected at the bradyzoite stage. Although there was a previous analysis that revealed the existence of TglncRNAs at *T. gondii* transcriptome from VEG strain^25^, with our novel approach it was possible to deepen the analysis and assign functions. LncRNA functional assignment is a complex task since the evolutionary relationship at sequence level is poor or undetected^6^. Obtaining concluding homology data from sequence is not possible in most cases, therefore, in our study we implemented at least three strategies in order to associate function to the predicted lncRNAS: 1) clustering algorithm over differentially expressed lncRNAs by coexpression with coding genes; 2) lncRNA genomic position relative to coding genes and 3) a *k*-mer profile analysis. We detected 217 differentially expressed TglncRNAs, process in *T. gondii*. Using a clustering algorithm we were able to group the 217 differentially expressed transcripts in 7 clusters, 5 of them integrated by TglncRNA. Interestingly, one of the clusters contains transcripts whose protein products are relevant for bradyzoite formation. One of them, TglncRNA-5J, being the most highly expressed TglncRNA at the bradyzoite stage. In addition, we were able to validate its differential expression in extracellular tachyzoites under stress conditions. Tachyzoite to bradyzoite stage conversion is a process that can be triggered by stress *in vitro*^47^. It is well documented that genes related with bradyzoite development, such as BAG1, can be induced by stressing extracellular tachyzoites^28^. Under this context it can be hypothesized that TglncRNA-5J might be involved in stress response and/or the differentiation processes regulation. In fact it is known in other model organisms that lncRNA expression is induced by stress^20^, while for parasites models organisms like *Trypanosoma brucei* a key regulation role of a new lncRNA named *gumpy* was documented during stumpy formation, a cell-cycle arrested stage induced by stress conditions^11^. Addtionally, in *P. falciparum*, the lncRNA-chr14 is a regulator of gametocyte formation^12^. In this sense, We consider that TglncRNA-5J is a good candidate for further experimental studies to know if it has a role in bradyzoite formation. Although the generation of clusters allowed us to determine possible associations of the TglncRNAs with co-expressed genes generating the possibility of association with biological processes and molecular functions, it is noteworthy that *T. gondii* ontologies are mapped from model eukaryotes losing essential information about some vital processes of the parasite like invasion, pathogenesis, differentiation, contingency genes. In accordance with this last observation, we explored other methodologies for associating function to TglncRNAs like the relative position of TglncRNAs loci to coding gene locations. The functionality of these lncRNAs might potentially be related to the regulation of transcription of these nearby genes in *cis*. There is a great amount of literature in the field that describe *cis*-lncRNAs activating gene expression when associated to regions demarcated as enhancers^48^. In this sense, our results revealed upregulated genes at the bradyzoite stage with loci close to TglncRNAs loci (i.e. BAG1), a characteristic that might indicate a potential function as enhancer-like^49,50^. On the other hand, the ROP2A gene which is downregulated, was found nearby to the TglncRNA-5J locus (upregulated). It is well known that *cis*-lncRNAs are able to repress gene expression by recruiting repressive protein complexes or by chromatin remodeling^51,52^. We highlight the observation of up and downregulated subtelomeric genes next to TglncRNAs loci. Specifically, one gene (*TGME*49_279460) of the multigenic family FamB was upregulated while a gene (*TGME*49_200130) of multigenic family FamC was downregulated. FamC and FamB genes were detected only in *Neospora caninum, Hammondia hammondi* and *T. gondii*^21^. Analysis in *T. gondii* indicates that they are subtelomeric genes, encoding integral membrane proteins with variable domains^53^. In addition, they can be also considered contingency genes, a feature that can be relevant for rapid response to environmental stress^53,54^. From the point of view of other parasites, the presence of these multigenic families FamB and FamC could be related to the multigenic family of subtelomeric *P. falciparum var* genes^55^. In this sense, it is interesting to note that *var* genes are regulated by *cis*-acting lncRNAs^36^. Given the heterochromatic nature of subtelomeres as well as the presence of few genes that may be associated with response to environmental stress, regulation by subtelomeric TglncRNAs could explain chromatin modulations specific in this chromosome domain in *T. gondii*. We also detected putative *trans*-acting TglncRNAs. Our analysis also shows possible interactions between TglncRNAs and upregulating transcripts at the bradyzoite stage like a DNA methyltransferase transcript (DNMTb)^56^. The lncRNAs role as mediators of DNA methylation at complex organisms has also been described^57^. Moreover, it was documented that DNMTs mRNA are stabilized by a RBP and a lncRNA mediated interaction, which conduct to more DNMT transcripts and more protein production^58,59^. Finally, we capitalize on the fact that positive correlated *k*-mer profiles between sequences allow the assignment of function, even when lncRNA from evolutionary distant organisms are compared^6^. We were able to find a related TglncRNA with the non-coding RNA activated by DNA damage, or NORAD *Homo sapiens* lncRNA that is relevant for genome stability in mammals^18,60^. Moreover, we observed Pumilio binding motifs at the predicted TglncRNA sequence that let us infer that the sequence would be functional^45^. Although Pumilio proteins were characterized in *T. gondii* and other parasites as regulators of translation by mRNA binding during differentiation processes^61–63^, no role at genome stability was proposed. In fact, a motif binding search for PUM proteins at *T. gondii* 5’-UTR and 3’-UTR transcripts reveals mRNA targets like those that codify for DNA repair proteins. In conclusion, our work proposes a bioinformatic approach for predicting TglncRNAs from transcriptomics data, in addition to functional association. Since our approach integrates a variety of scalable bioinformatics tools, it would be possible to apply it in other protozoans and non-model organisms with available transcriptomics. Furthermore, we believe that other data sets like ribosome profile, if available, could be integrated to validate non-coding transcripts. At this point it is worth mentioning that a ribosome profile data set is available for *T. gondii* only at the tachyzoite stage which makes it not adequate for the bradyzoite context presented in this work^64^. Finally, we believe that although empiric validation is needed to confirm our observations, this work brings a useful perspective to amplify lncRNAs study in the protozoan pathogens field.

## Methods

### Data acquisition

Raw sequence data for tachyzoite and bradyzoite stages were obtained from^22^. Genome sequences, transcripts UTRs and feature files for *Toxoplasma gondii*, strain ME49 and VEG, were obtained from ToxoDB v59. TglncRNA sequences from VEG strain were downloaded from the RNAcentral database (RNAcentral.org^65^,release 22). All gene ontologies enrichment analysis were performed at ToxoDB.

### RNA seq analysis

Raw sequences were evaluated by FASTQC software (https://github.com/s-andrews/FastQC). Adaptors and quality trimming were performed by TrimGalore! software (https://github.com/FelixKrueger/TrimGalore). Surviving reads were mapped against the T. gondii genome (v59). Assembly was performed by StringTie protocol^66^. Statistical analysis was performed by the DESeq2 package^67^. Transcripts with *log*_2_ *f c >*= 2 & *log*_2_ *f c <*= −2 and *p*−*ad justedvalue <* 0.05 were considered for further analysis. Volcano plots were made using the Enhanced Volcano package (https://github.com/kevinblighe/EnhancedVolcano). Heatmaps were made using the Complex Heatmap package (10.18129/B9. bioc.ComplexHeatmap). Clustering was performed by the kmeans algorithm with the following parameters: nº of centers 7, maximum number of iterations 1000 and 10 random sets. The optimal *k*-value was founded by the Elbow Method.

### TglncRNA prediction and analysis

The bioinformatic approach is resumed at Fig. 1a. Annotated transcripts were filtered from transcriptome assembly, that include coding and non coding transcripts (rRNA and tRNA). A custom bash script was employed to filter out unannotated transcripts based on sequence length > 200 nt. These transcripts were then analyzed for their coding potential using the Coding Potential Calculator Tool (CPC2^68^). Transcripts classified as “non-coding” by CPC2 software were further examined using the blastx tool^69^ against SwissProt databases^23^. Expressed sequence tags for *T. gondii* (v59) were downloaded from ToxoDB and predicted sequences for lncRNAs were aligned against ESTs using blastn. The best match for each EST was considered (*E* −*value <* 1*e*− 05). Relative expression for coding and predicted TglncRNA transcripts (*f pkm*) was obtained by the ballgown package^66^. A Mann–Whitney U test was performed to compare means of relative expression between tachyzoite and bradyzoite coding and TglncRNAs transcripts (*p*-value < 0.01). GC content, sequence length distributions and plots were calculated and made by a custom python script. Chromosome plots for predicted TglncRNA loci were made with chromoMap^70^. Analysis of TglncRNA loci distances to gene and plots was made with ChiPSeekr^71^, distances to TSS was set to +*/* − 10*kbp*. TglncRNA homologous between strains (ME49 vs VEG) were determined by a blastn analysis (*E* −*value <* 0.001). Venn diagram was constructed using a web tool (https://bioinformatics.psb.ugent.be/webtools/Venn/). Circos plot was made with the circlize package^72^. TglncRNAs loci at VEG chromosomes were determined by a blastn analysis (*E*-value < 1*e*− 10). Coverage plots were made using the pyGenomeviz package (https://github.com/moshi4/pyGenomeViz). Coverage files (BigWig) were created using bamCoverage tool (default parameters except for normalization: CPM) from deeptools package^73^. To predict TglncRNA interaction with coding transcripts, RIBlast^41^ software was employed using the following parameters: MaximalSpan:70, MinAccessibleLength:5, MaxSeedLength:20, InteractionEnergyThreshold:-4, HybridEnergyThreshold:-6, FinalThreshold:-20, DropOutLengthWoGap:5, DropOutLengthWGap:16. Only differentially expressed transcripts at the bradyzoite stage were considered for the analysis.

### Network analysis

Pearson’s correlations between differentially expressed transcripts (coding vs TglncRNAs) at bradyzoite stage were calculated taking into account relative expressions (*log*_2_ *f pkm*). Matrix was filtered by *r*-values > 0.6 & *p*-value < 0.01. Only correlations between coding transcripts and the predicted TlgncRNAs by RIBlast were considered. Network was analyzed and plotted using the igraph package (https://igraph.org). Communities were calculated using the Louvain algorithm.

### *K*-mer profile analysis

The predicted *T. gondii* lncRNAs were analyzed using the package seekr^6^. To avoid bias only the first isoform per TglncRNA was considered in the analysis. For *k*-mer count the module seekr_kmer_counts was used setting *k*-size from *k* − 5 to *k* − 7. In each case no matrix with null elements was obtained. For prediction of RNA binding protein (RBP) motifs the module seekr_pwm was employed. As input we used the full data set of position weighted matrix (PWM) from CISBP-RNAdb^43^ and the 3 above calculated matrix of *k*-mer profiles (*k* − 5,*k* − 6 and *k* − 7). Three weight matrices of TglncRNA vs PWM motifs (pwd_sum) were obtained, *log*_2_ transformed and used for further analysis. The *k* − 7 weighted matrix was filtered by *log*_2_(*pwd*_*s*_*um*) *>* 6. Finally, for each TglncRNA in the above filtering matrix, the 10 highest weights for each motif were kept for further analysis.

The canonical RBP for the identified motifs were obtained from CISBP-RNAdb and *T. gondii* homologous were determined by the interpro domain tool at ToxoDB (Supplementary Table S5). For the *k*-mer profile comparative analysis, the *Homo sapiens* lncRNAs for the canonical RBP were obtained from RAIDdb^44^. Corresponding sequences were obtained from RNAcentral.org: ANCR (*URS*0000*A*4*F*650_9606), MALAT1 (*URS*0000*A*77400_9606), CRNDE (*URS*0002123222_9606), BCYRN1 (*URS*0000193*C*7*E*_9606), NEAT1 (*URS*000075*DAEC*_9606), NORAD (*URS*000075*CEFB*_9606), HOTAIR (*URS*000075*C*808_9606), XIST (*URS*000075*D*95*B*_9606). The input data set was the 10 TglncRNAs with best PWM weight for each motif obtained previously. Additionally all the first isoforms sequences for the known *H. sapiens* lncRNAs were obtained using the module seekr_download_gencode (GENCODE^74^,v44). These sequences were employed to generate a normalization vector using module seekr_norm_vectors. A *k* − 5 was used as a parameter since no null elements were generated in the reference. *K*-mers count (*k* − 5) was performed for the reference set of *H. sapiens* and the input *T. gondii* dataset using the normalization vectors calculated above. Pearson correlations were calculated using module seekr_pearson. The *r*-values Pearson matrix was filtered by *r*-value > 0.2 & *p*-value > 0.01. Dot plot analysis was performed at https://fasta.bioch.virginia.edu/fasta_www2/fasta_list2.shtml using the *plaling* tool. Multiple sequence alignment was made using the MAFFT algorithm at JalView^75^ suite.

### *In vitro* stress followed by RT-qPCR assay

Freshly extracellular tachyzoites of RHΔhxgprt strain were incubated 2 hrs at 37^*°*^*C* (unstressed,Tz) or 43^*°*^*C* (stressed, St). After that they were conserved in TriZol (Invitrogen) solution at −80^*°*^*C* until use. In both cases, each experiment was performed in three independent biological replicates. RNA extraction was performed according to the manufacturer’s instructions and cDNA was obtained by means of MMLV reverse transcriptase (Promega) using oligo dT with the protocol provided with the enzyme. For RT-qPCR primers used are listed in Supplementary Table S7. The primers were first assayed for efficiency, and the threshold was defined for each set of primers. Melt curves were also obtained for each set. For each set of experiments, SybrGreen master solution (Roche) was used and qPCR was run in StepOne Real time equipment (Applied Biosystems). Actin and tubulin were used as housekeeping genes and data was normalized to those genes by Linereg^76^ free software (v2021.2), applying the geometrical median. All sets of data were relativized to tachyzoite amplification.

## Acknowledgements (not compulsory)

We thank our colleagues in the lab for helpful discussions. This research was supported by a Grant from the Agencia Nacional de Promoción Científica y Tecnológica (ANPCyT) Grant BID PICT 2019-0513. L.V., S.O.A. and A.M.A are researchers from the National Council of Research (CONICET) and UNSAM. The funders had no role in study design, data collection and analysis, decision to publish, or preparation of the manuscript.

## Author contributions statement

Conceived and designed the experiments: A.M.A., L.V. and S.O.A. Performed the experiments: L.V.,G.L.,C.C and A.G. Analyzed the data: A.M.A. and L.V. Wrote the paper: A.M.A., L.V.,C.C. and S.O.A. All the authors were involved in reviewing and editing the manuscript. All authors contributed to the article and approved the submitted version.

## Data availability statement

Public raw data used in this work is available at https://ncbi.nlm.nih.gov/geo/query/acc.cgi?acc=GSE132248.

