## Supplementary Data for "A bioinformatic approach for the prediction and functional classification of *Toxoplasma gondii* long non-coding RNAs"

[2] Escuela de Bio y Nanotecnologías (UNSAM)

### 1 Legends to Supplementary Tables

**Supplementary Table S1:** Primers used in this work. In the table are listed all the primers used in RT-qPCR assays, with the forward and reverse sequence, product size and average efficiency (three independent assays). Gene IDs: Tgtubulin: $TGME49_266960$; Tgactin: $TGME49_209030$; Tgsag1: $TGME49_233460$; Tgbag1: $TGME49_259020$.
**Supplementary Table S2:** TglncRNAs genome localization. In the table are listed the lncRNAs found in the analysis. lncRNA: *T. gondii* lncRNA ID; chromosome: name of TGME49 chromosome; start and end: position of a lncRNA locus at the chromosome; Bradyzoite and Tachyzoite: $log_2 fpkm$ values for the corresponding TglncRNA transcript; Uniprot hit: results from blastx experiment between TglncRNAs vs SwissProt database. Bold highlighted IDs correspond to TglncRNAs analyzed by qPCR in this work.
**Supplementary Table S3:** Differentially expressed TglncRNAs between tachyzoite and bradyzoite stages. Transcript Id: id assigned to the predicted lncRNA transcript by the StringTie assembler; $p-value$: Wald test $p-value$; padj: Benjamini-Hochberg adjusted $p-value$. Transcripts analyzed in this work are highlighted in bold typography. Bold highlighted IDs correspond to TglncRNAs analyzed by qPCR in this work.
**Supplementary Table S4:** Gene ontologies (GO) enrichment analysis. Table is organized in 7 sheets each one corresponding to one cluster. Each sheet is organized as follows. Transcripts id: ids for each transcript that composes the cluster; TglncRNA: lncRNAs found in the cluster. Molecular and Biological ontologies are organized in two tables that share the next columns; ID: GOs identifiers; Name: GO descriptions; Bgd count: number of GO in background proteome (TGME49v59); Result count: number of GO in the sample; Result gene list: genes associated with the corresponding GO in the sample; Pct of bd: percent of GOs in background proteome; Fold enrichment: enrichment of GO in sample relative to background proteome; Odds ratio: Odds ratio static from Fisher's exact test ; P-value: $p-value$ from Fisher's exact test; Benjamini and Bonferroni: adjusted $p-values$.
**Supplementary Table S5:** Cis- and trans-acting Functional Classification of TglncRNAs. Table is organized in 2 sheets. 1-Predicted cis-TglncRNAs: 46 genes differentially expressed and closer to at least one TglncRNA locus; Gene id: identifiers for each gene; Description: gene product description. Subtelomeric genes (ST) are highlighted; log2FoldChange: transformed logarithmic of relative ratio of change for each gene; padj: Benjamini and Bonferroni adjusted p-values. TglncRNA: closer (less than 10 kbp) locus to a coding gene. LncRNA analyzed in this work are highlighted in bold typography. 2-Predicted trans-TglncRNAs: TglncRNA and 117 transcripts predicted as interaction targets. Transcripts id: identifiers for each coding transcript; Description: product description for each coding transcript; TglncRNA: lncRNA predicted by RIBlast to interact to the corresponding coding transcript; Transcript community: number of community that a coding transcript integrates in the correlation network from Figure 5C; TglncRNA community: number of community that each TglncRNA from column "TglncRNA" integrates. Bold highlighted IDs correspond to TglncRNA analyzed by qPCR in this work. Bold highlighted IDs correspond to TglncRNAs analyzed by qPCR in this work.
**Supplementary Table S6:** Predicted RNA binding proteins for TglncRNAs. File is composed by two sheets: 1-Predicted RNA binding proteins (RBP); Motif ID: RNA motif identifiers; CISBP-RNA ID: database RBP identifier; Name: RBP common name; Description: RBP gene description; Gene ID: cross-referenced databases gene identifiers; Interpro Id: interpro domain identifiers; Domain Name: interpro domain description;

Homologous in Tg: homologous RBP at *T. gondii* proteome (v59); Target TglncRNA: predicted TglncRNA targets for each RBP. The 10 sequences with highest weight for each motif were considered for the analysis. The analyzed sequences in this work are highlighted in bold typography. 2- HslncRNA correlations. Pearson correlations ($r - value$) for k-mer profiles between TglncRNAs predicted as targets for a RBPs and validated Homo sapiens lncRNAs as targets for the homologous RBPs. TglncRNA: lncRNA sequence used for the analysis (the analyzed sequence in this work is highlighted); RBP: canonical predicted RNA binding protein from CISBP-RNA database; HslncRNA & HsRBP: *H. sapiens* lncRNA and the validated RBP for each RNA sequence according RNA Association Interaction Database (RAID); $r - value$: pearson's correlation values; $p - value$: statics for pearson tests. Bold highlighted IDs correspond to TglncRNAs analyzed by qPCR in this work.

**Supplementary Table S7:** Predicted PUM targets GO enrichment analysis represented by a word cloud. Table summarize the 14 genes related with DNA damage repair and metabolism process and is organized as follows: Gene Id: identifier from ToxoDb; Product Description: description about gene product; Computed GO process IDs: biological process identificators; Computed GO Processes: biological process description; PRE: 5'-UTR sequence region where PUM binding motif (TGTANATA) was found. for the GO enrichment analysis of predicted PUM targets. DNA maintenance processes are highlighted with a red dashed box line.

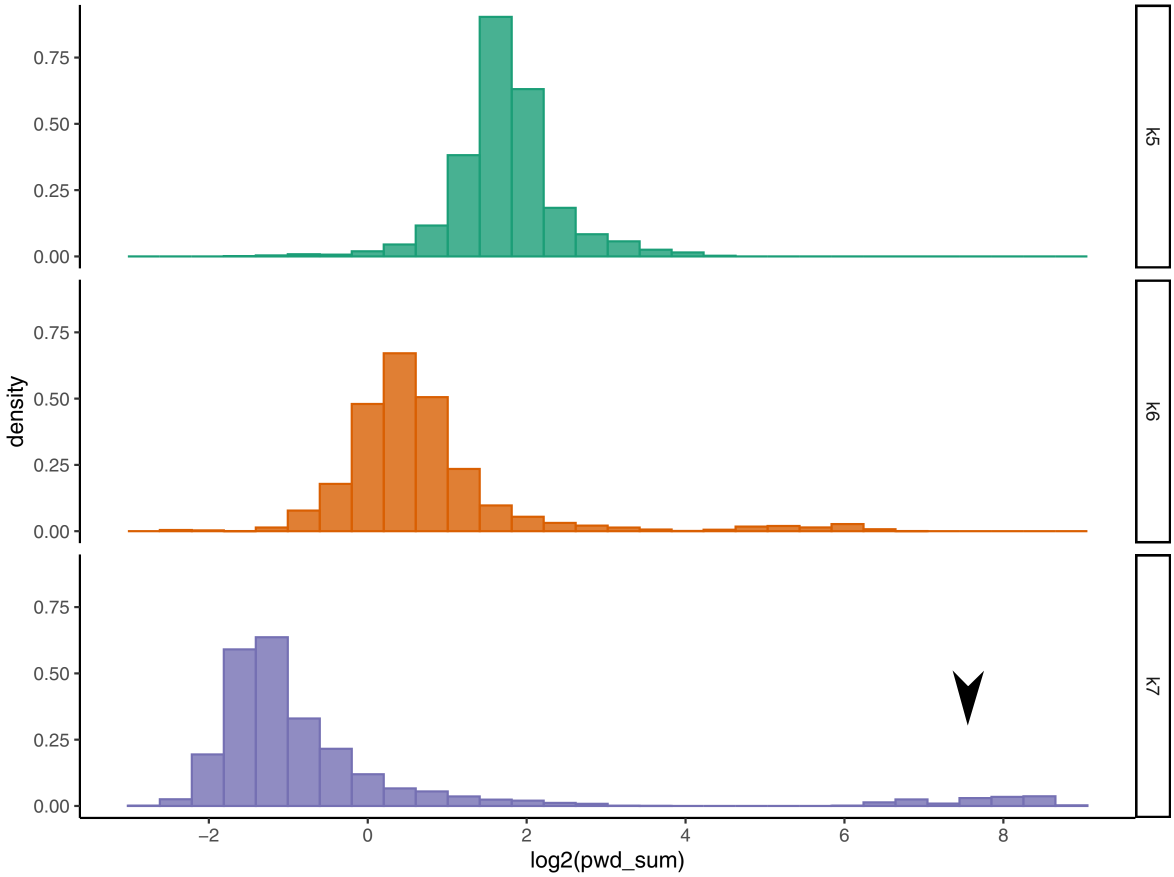


**Supplementary Figure S1:** Density distribution of PWM weights for $k-5$, $k-6$ and $k-7$. High weighted PWM at $k-7$ distributions is highlighted by an arrow. $log_2(pwm\_sum)$: transformed weight for a PWM motif.

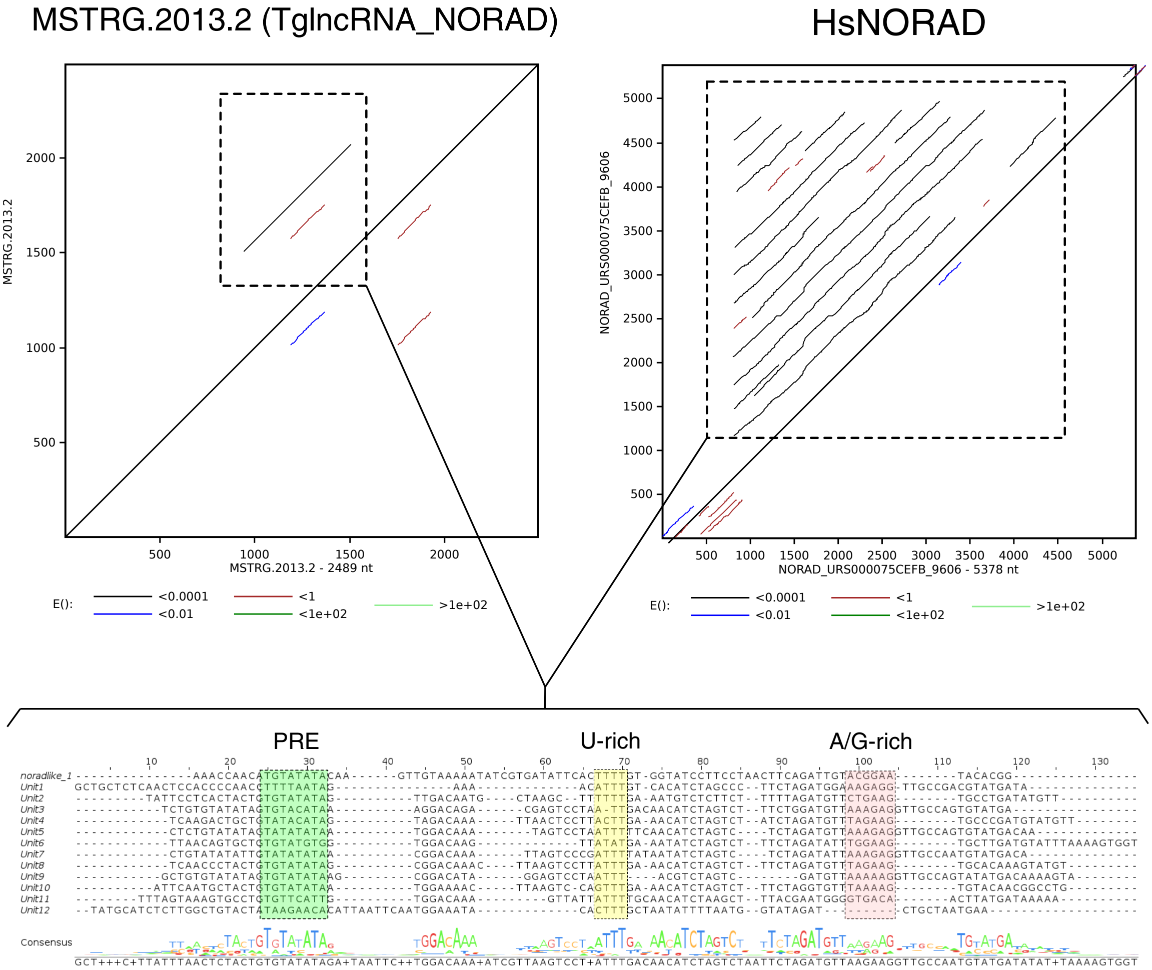


**Supplementary Figure S2:** Dot plot analysis for the predicted TgNORAD-like lncRNA ($TglncRNA\_NORAD$). Upper panel: Dot plot analysis for $MSTRG$.2013.2 ($TglncRNA\_NORAD$) and *Homo sapiens* NORAD ($URS$000075$CEFB_$9606) . Black line diagonal represents full identity, a black line over diagonal represents a sequence repetition with high significance ($E - value < 0.0001$). Bottom panel: multiple sequence alignment (MSA) between 1 repetition ($noradlike\_1$) in $MSTRG$.2013.2 and the 12 documented units of repetitions in the human NORAD ($Unit1 - 12$). Conserved motifs are highlighted .PRE: Pumilio recognition elements.
